# 3D organoids containing endothelial and neural cells generation by serial inductions of differentiation on human iPSC-derived embryoid bodies

**DOI:** 10.1101/2025.05.20.653559

**Authors:** Tongguang Wang, Anna Bagnell, Valerie McDonald, Benjamin D. Gastfriend, Joseph P. Steiner, Abdel G Elkahloun, Kory Johnson, Rebekah G. Langston, Mark R. Cookson, Avindra Nath

**Author notes:** Correspondence should be addressed to: Dr. Tongguang Wang; Dr. Avindra Nath.

## Abstract

3D brain organoids have been widely used as a tool to study human brain development and disorders. Although endothelial cells play important roles in the brain development and pathogenesis in neurological disorders, most 3D brain organoids lack inherent endothelial cells and need either the addition of endothelial cells or to be transplanted to animals to reconstitute such vascular structures, likely missing the developmental interactions of endothelial cells and other cells in the human brain. In order to reconstitute a 3D organoid mimicking the in vivo neural and endothelial cells development, we cultured iPSC-derived embryoid bodies in sequentially applied endothelial and neuronal induction media along with Matrigel embedding. The resulting 3D organoid consists of both neural cells and endothelial cells with vascular like structures, as determined by immunostaining. With scRNA-Seq analysis, the brain organoid was confirmed to contain neural cell types similar with human brains, including a variety of excitatory and inhibitory neurons and glia. Furthermore, when compared with traditional cerebral organoids without endothelial cells using RNA-Seq analysis, the endothelial containing neural organoids (EC-neural organoids) showed difference in gene profiles and favored angiogenesis and vasculogenesis. Of the differentially expressed genes, KRBA2 expression was found higher in neural cells and its inhibition by siRNA treatment resulted in decreased transcriptions of a variety of genes such as neuronal differentiation specific genes but not in genes specific to pluripotent stem cells such as OCT4. The EC-neural organoids also express receptors to SARS-CoV-2 similar to human brains. This 3D model provides a useful tool to study the interactions of endothelial cells and neural cells in the brain development and neural infectious disorders where endothelial cells and pericytes play pivotal roles.

## INTRODUCTION

Organoids derived from human stem cells have been developed in recent years as an alternative in vitro model for animal and human subject studies. With relatively less ethical concerns and wide feasibility, human organoids constitute multiple cell types and tissue structures similar to the corresponding human organs. Set to partly substitute animal models, they represent the future of human disease modeling and have the potential to play a pivotal role in the study of disease pathogenesis, drug development and for guidance of individualized therapies, as indicated by the reported successes in the treatments of cystic fibrosis (1)and some tumors. As such, studies using 3D brain organoids derived from human induced pluripotent stem cells (iPSCs) have also prevailed in the field of neuroscience research, modeling neural infectious diseases including ZIKA, SARS-CoV-2 and neurodegenerative diseases including amyotrophic lateral sclerosis (ALS) and Alzheimer’s disease. However, conventional 3D neural organoids lack certain cell types, such as microglia and endothelial cells as they were derived from other germ layers than the other neural cells. While evidences suggest that the pathogenesis of many neurological disorders is not limited to neurons or central nervous system but are the result of complicated interactions involving multiple organs, even from imbalances in the gut microbiota(2). As for modeling neural infectious diseases, it is inadequate to use the neural organoids without endothelial cells and microglia, as a lot of infectious agents, such as SARS-CoV-2, must overcome the blood brain barriers to access the neural cells before infecting the neural cells. Without endothelial cells, the model will be incomplete and may fail to provide clinically relevant knowledge. To reconstitute endothelial or blood vessel into the brain organoids, differentiated endothelial cells have been added separately to neural spheres(3). Alternatively, the neural organoids can be transplanted into an animal brain to get vascularized(4). These approaches either missed the initial interactions of endothelial cells and neural cells during development or involved complicated techniques such as animal operations which undermined the benefit of using in vitro 3D brain organoids.

The interactions between neural cells and endothelial cells play important roles in the development of neural system and its normal function. Besides conduits for the delivery of oxygen and nutrients, perfusion independent roles of organotypic endothelium play during brain development and regeneration through releasing specific paracrine growth factors called angiocrine growth factors or angiokines(5). For example, we have reported that VEGF can function as a regulator for oligodendrocyte proliferation and differentiation(6). On the other hand, it is well known that endothelial cells exhibit distinct functional and molecular heterogeneity depending on the type of vessel, organ and especially age. Although it is not yet well studied, it is conceived that endothelial cells and hematopoietic cells develop in parallel and the later arise from specialized endothelial cells(7). There are endothelial cells that exist as stem/progenitor like cells which continuedly proliferate and are ready for angiogenesis even in adults. It is known that different from neural cells originated from exoderm, endothelial cells are from mesoderm whose specification is initiated by BMP4 and bFGF(8). Expression of VEGFR2 on angioblasts is necessary for allowing their communication with the binding ligand VEGF-A which further facilitate endothelial cell development in a highly controlled manner. Thus, it is possible to derive the endothelial cells by mimicking the in vivo process along side with the neural differentiation.

To address these issues, we developed a 3D brain organoid model by inducing the endothelial cell differentiation with serial applications of BMP4, bFGF and VEGF to iPSC-derived embryoid bodies, then with neuronal differentiation differentiation and maturation media. The resulting organoids contain endothelial cells and neural cells similar to human brains. By comparing them with traditional neural organoids, we found certain differentially expressed genes which may play unique roles in the neural development. Besides, we screened for SARS-CoV-2 receptors on the endothelial containing organoids (EC-neural organoids) and found the similar distribution as in human brains. These results indicate the EC-neural organoids can be used as a model to study human neural development and potentially, neural infections such as SARS-CoV-2.

## MATERIALS AND METHODS

### Reagents and supplies

Culture media and components were purchased from Invitrogen (Carlsbad, CA); growth factors and cytokines were purchased from PeproTech (Rocky Hill, NJ) and chemicals were purchased from Sigma (St. Louis, MO) if not otherwise specified. Details of the reagents and resources were provided in the SI Appendix, Table S1.

### Cell cultures

Blood from adult healthy donors was collected at the Transfusion Medicine Blood Bank of the Clinical Center at the National Institutes of Health (NIH). Signed informed consents were obtained in accordance with the NIH Institutional Review Board. Human induced pluripotent stem cell (iPSC) lines 505, 506 and 516 were generated from PBMCs enriched from the blood samples using Sendai virus method by National Heart Lung and Blood Institute iPSC core facility at NIH as published previously(9). The iPSCs were cultured in E8 Flex medium on Matrigel coated plates in a 5% CO_2_ incubator at 37⁰C and sub-cultured using ethylene diamine tetra acetic acid (EDTA), following published protocols(10).

### Culture 3D brain organoids containing endothelial cells

Embryoid bodies were derived from iPSCs using AggreWell™ plates (Stemcell Technologies, Canada) according to manufacturer’s instruction. Briefly, iPSCs were dissociated into single cells by treatment with Accutase dissociation buffer (Invitrogen) and counted. One million cells per well were seeded onto a low attachment 24-well AggreWell plate in E8 Flex medium with Rock inhibitor Y-27632 (10 μM). After 24 hours, embryoid bodies were formed and transferred to new 24 well plates pretreated with anti-adherence rinsing solution (Stemcell Technologies) at ~10 spheres per well in 1 ml of E8 Flex medium.

To form 3D organoids consisting of neural and endothelial cells, 48 hours after being transferred to a new plate, the embryoid bodies were coated by adding ice-cold Matrigel drop wisely (25 ul/500 ul medium) into the medium and then incubated in a 5% CO_2_ incubator at 37⁰C for 4 hours. The endothelial cell induction process was done following modifications to a protocol that used in 2D endothelial induction from iPSCs (11). Briefly, 500 ml of medium consisting of DMEM/F12, 1X B27 supplements and Activin A (125 ng/ml) was added on top of the spheres followed by incubation for 24 hours. Then 500 ml medium was replaced with freshly made E8 Flex consisting of 10 ng/ml BMP4 followed by another 25 ul of ice-cold Matrigel overlay and incubation for 72 hours. The medium was then changed with DMEM/F12 consisting of 1X B27 supplement, 1 mM 8-bromo-cAMP, 100 ng/ml vascular endothelial growth factor (VEGF) and antibiotics for 72 hours. After that, half medium was changed with DMEM/F12 medium consisting of 1X N2 supplements, 1X B27 supplements, 20 ng/ml VEGF and antibiotic-antimycotic was done every other day to facilitate neural differentiation and maintain the growth of the organoids. The organoids were transferred to a 6 well plate pretreated with anti-adherence rising solution after 2 weeks in culture to avoid over-crowding of cells.

3D cerebral organoids without endothelial cell induction were also derived from iPSCs using the Kit from Stem Cell technologies with modifications. Following the embryoid body formation using the Aggrewells, the neural induction and expansion were done as instructed and at day 5, the Matrigel was used to coat the spheroids drop wisely. The organoids were then changed to maturation media for further neuronal maturation.

### 3D Organoid Clearance and Immunocytochemistry

For immunostaining, a clearing process was performed on the organoids using a Kit purchased from Visikol, following the manufacturer’s instruction with optimization. Briefly, the organoids were fixed in 4% paraformaldehyde (PFA) for 24 hours in a cold room and then washed 2 times, 1 hour each, with phosphate buffered saline pH 7.4 (DPBS) to remove the PFA residue. Then dehydration/permeabilization was done at 4°C with gentle shaking for 10 mins in each of the following buffers subsequentially: 50% methanol in PBS, 80% methanol in water, 100% methanol, 20% DMSO in methanol, 80% methanol in water, 50% methanol in PBS, 100% PBS and PBS with 0.2% triton X-100. After incubation for 30 minutes with penetration buffer (0.2% triton X-100, 0.3M glycine and 20% DMSO), the organoids were blocked with blocking buffer (DPBS with 0.2% triton X-100, 6% donkey serum and 10% DMSO) for 30 mins. The organoids were then incubated with the primary antibodies (rabbit anti-MAP2, Abcam; rabbit anti-NG2, Abcam, rabbit anti-PDGFR-beat and mouse monoclonal anti-CD31, Thermo Fisher Scientific) at 1:100 in DPBS with 0.2% Tween 20, 3% donkey serum and 5% DMSO for 1 hour at 37°C and then followed by 4°C overnight. After that, the organoids were washed five times in DPBS with 0.2% Tween 20 and 100 ug/ml heparin for 10 minutes each time. The organoids were then incubated with corresponding secondary antibodies (1:100, anti-mouse Alexa Fluor 488: Thermo Fisher Scientific Cat# A-11001; anti-rabbit Alexa Fluor 594: Thermo Fisher Scientific Cat# A-11012, RRID:AB_2534079) at 37°C for 1 hour followed by 1 hour of 2-(4-amidinophenyl)-1H-indole-6-carboxamidine (DAPI) nuclear staining at room temperature. Then the organoids were washed for at least 10 times at 37°C with shaking, 10 minutes each. When ready for imaging, the buffer was replaced with 200 ml of HISTO M clearing solution (Visikol). Images were acquired using a Zeiss LSM 510 META multiphoton confocal system (Carl Zeiss) or ImageXpress Micro confocal microscope (Molecular Devices).

### Single cell RNA-Seq analysis

Organoids were dissociated to single cells by treatment with Accutase for 20 minutes with shaking. Single cell suspensions quality, number and viability were assessed with a dual fluorescence cell counter Luna-FL (Logos Biosystems). 8,000-10,000 cells were targeted from each normal cell suspension. The cells were washed twice with PBS+0.04% bovine serum albumin and resuspended at about 1,000 cells per microliter. 10X Genomics’s Chromium instrument and Single Cell 3⍰ Reagent kit (V3 and 3.1) were used to prepare the individually barcoded single-cell RNA-sequencing libraries following the manufacturer’s protocol. Quality of the libraries was assessed by the Bioanalyzer traces (Agilent BioAnalyzer High Sensitivity Kit) and quantitated by the Qubit system. Sequencing was done on the Illumina NextSeq machine, using the 150-cycle High Output kit with 28⍰bp read 1, 8⍰bp sample index, and 91⍰bp read 2. Following sequencing, the bcl files were demultiplexed into a FASTQ, aligned to human transcriptome GRCh38 and single-cell 3⍰ gene counting was performed by the standard 10X Genomics’s CellRanger mkfastq software (V3.0.2). The gene expressions in single cells were visualized using 10X Genomics Loupe Browser 4.0.0. Pseud time of differentiation Trajectories was annotated by using Partek flow scRNA-Seq analysis tools. (https://partekflow.cit.nih.gov).

### Neural stem cell differentiation and small inhibitory RNA (siRNA) transfection

Neural stem cells (NSC) were derived directly from cord blood CD34 cells (CD34-iNSC)(12) or differentiated from iPSCs (NSC 507 and NCRM) as reported in our previous publication(13). For differentiation from iPSCs, iPSCs were seeded on a Matrigel-coated 6 well plate in E8 medium. When ready for induction, the E8 medium was replaced with PSC neural induction media (Gibco) for 7 days. The resulting NSC were characterized by nestin immunostaining and were further differentiated to neurons when subcultured on Matrigel-coated plates and incubated with neuronal differentiation medium (DMEM/F12 containing 1xN2 supplement, 1xB27 supplement, 300 ng/ml cyclic adenosine monophosphate (cAMP, Sigma) and 0.2 mM vitamin C (Sigma), 10 ng/ml brain derived neurotrophic factor (BDNF) and 10 ng/ml glial-derived neurotrophic factor (GDNF) for 14 days. The resulting neurons were confirmed by immunostaining for β-III-tubulin. Premade TriFECTa RNAi kit targeting human KRBA2 (hs.Ri.KRBA2.13) and IGFBP6 (hs.Ri.IGFBP6.13.1, hs.Ri.IGFBP6.13.2 and hs.Ri.IGFBP6.13.3) as well as the non-specific control RNAi were purchased from IDT. For siRNA transfection, NSCs were cultured for 24 hours and reached 80% confluence. TransIT-siQUEST transfection reagent (Mirus Bio, Madison, WI, USA) was mixed with RNAi (30 nM final concentration). After 15 min, the mixture was applied to the NSC cultures. The cells were then lysed for RNA purification after 48 hours.

### Real-time polymerase chain reaction (RT-PCR) assay

RNA was extracted and purified using a total RNA PLUS purification kit (Norgen) following manufacturer’s instructions. First strand cDNA was synthesized using QuantiTect Reverse Transcription kit (Qiagen) following the manufacturer’s instruction. qPCR was conducted using FastStart Universal SYBR Green Master (ROX) kit from Roche and ViiA7 real-time PCR system (Life Technologies) according to manufacturer’s instructions. Herv-K Env, Nestin, MAP2 and beta-2-microglobin (B2M) primers were manufactured by Invitrogen as published before(14). KRBA2 and IGFBP6 primers were purchased from IDT. The sequences of the primers were reported in the supplemental Table 1. Delta Ct Value (ΔCt) was calculated using the accompanying software with the ViiA7 real-time PCR machine. ΔΔCt method was used for the differential gene expression analysis.

### Analysis of published literature

Allen Atlas’s microarray database (2010 Allen Institute for Brain Science. Allen Human Brain Atlas. Available from: human.brain-map.org)(15) were used to compare the gene expression profiles between brain organoids and normal human brains. FURIN and BSG in human brains. The cell specificity of gene expression was checked using cell transcriptomic RNA-Seq data tool in the Allen brain database (Allen Cell Types Database 2015, http://celltypes.brain-map.org/rnaseq/human_m1_10x). These results are shown as heatmaps. The levels of ACE2 and TMPRSS2 were also examined in a developing brain (16) using the UCSC cell browser v0.7.11(http://genome.ucsc.edu/)(17).

### Experimental design and statistical analysis

For conceptual proving, three batches of brain organoids from different iPSC lines were derived independently and immunostained for the neuronal and endothelial cell markers. Four brain organoids were used to study the levels of SARS-CoV-2 infection associated genes using scRNA-Seq analysis. Three were also used for their distributions among cell types. For statistical analysis, at least three independent experiments were performed. Mean and standard error (SE) were calculated for each treatment group. For RT-PCR results, ΔCt or ΔΔCt were calculated. For comparison of gene expression, the levels of gene expression were presented as fold of the highest expression treatment group. As a result, one sample t test was performed against the hypothetical value 1. Statistical analysis was performed using Prism, version 3.0.

### Data availability

All relevant data supporting the key findings of this study are available within the article and its Supplementary Information files. The RNA-Seq data are deposited in public domain.

### Code availability

Softwares used in this study include 10X Genomics Loupe Browser 4.0.0 (https://support.10xgenomics.com/single-cell-gene-expression/software/visualization/latest/installation)

## RESULTS

### 3D neural organoids generated contain neural and endothelial cells

To generate a neural organoid similar to human brain, which consists of neurons, glia and endothelial cells, we developed a protocol to first differentiate iPSC-derived embryoid body partially to endothelial cells then followed with incubation in medium supporting both endothelial and neural cells. By using AggreWells, the uniform sized embryoid bodies were generated from iPSCs. After 48 hours, Activin-A was applied into the medium following Matrigel coating of the embryoid bodies. 24 hours later, media was replaced with DMEM/F12 media containing BMP4 and bFGF to initiate the endothelial cell differentiation. At day 6, the media containing cAMP and VEGF was used to finalize the endothelial differentiation and initiate the neuronal differentiation. After 72 hours, the resulting organoids were further cultured in media containing N2, B27 supplements and VEGF to support both neural and endothelial cells (Fig 1A). Shortly at this stage, the organoids showed a cyst-like structure, consisting of tightly connected monolayer of cells, these cells then developed into CD31+ endothelial cells (Fig 1B). A mass core then appeared within the organoids, where most MAP2+ neurons were located. As seen in Fig 1C, after clearance and immunostaining, at 8 weeks after differentiation, the organoids contained both MAP2+ neuronal cells and CD31+ endothelial cells. At this stage, the endothelial cells were mainly lining the organoids while neuronal cells occupying the inner mass. When further cultured until 12 weeks, the CD31+ endothelial cells were seen within the organoids, showing blood vessel like structures (supplemental Z-scan movie and Fig 2C). To further support that vascularization existence in the organoids, pericyte markers PDGFR-beta and NG2 were coimmunostained with CD31. Both pericyte markers showed certain degrees of colocalization with CD31 (Fig 2A). The resulting organoids also contained few microglial cells, as shown by the TREM2+ staining cells in the organoids (Fig 2B).

**Figure 1.**
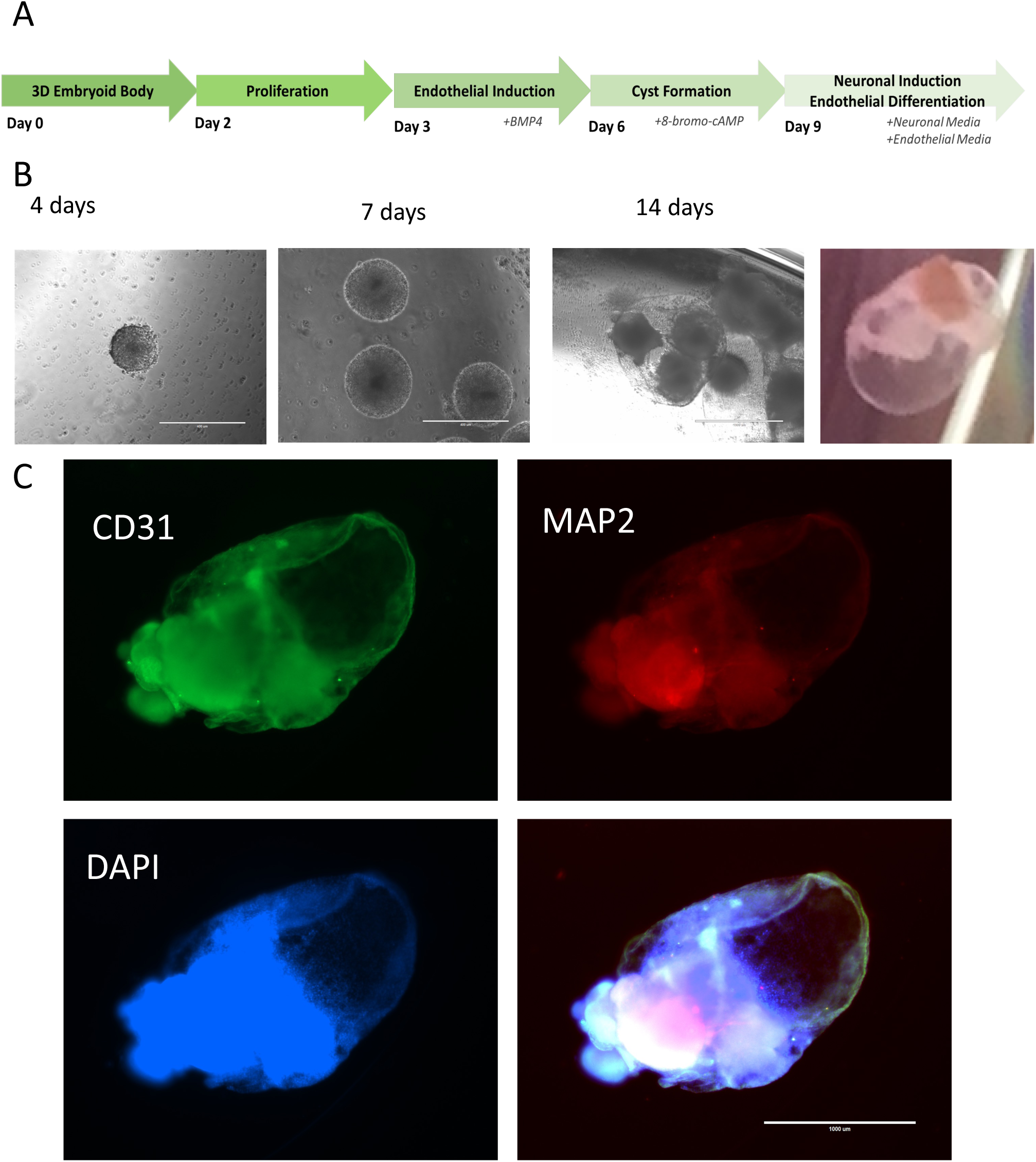
Derive 3D neural organoids with endothelial cells. 3D organoids were derived from iPSCs following the schematic procedures to achieve endothelial cells and neural cells simultaneously (A). The morphological changes through the organoid development (B). The resulting organoids at age of 8 weeks were immunostained and showed CD31+ endothelial cells lining over a mass enriched with MAP2+ neurons (C). Representative images from three independent experiments were shown.

**Figure 2.**
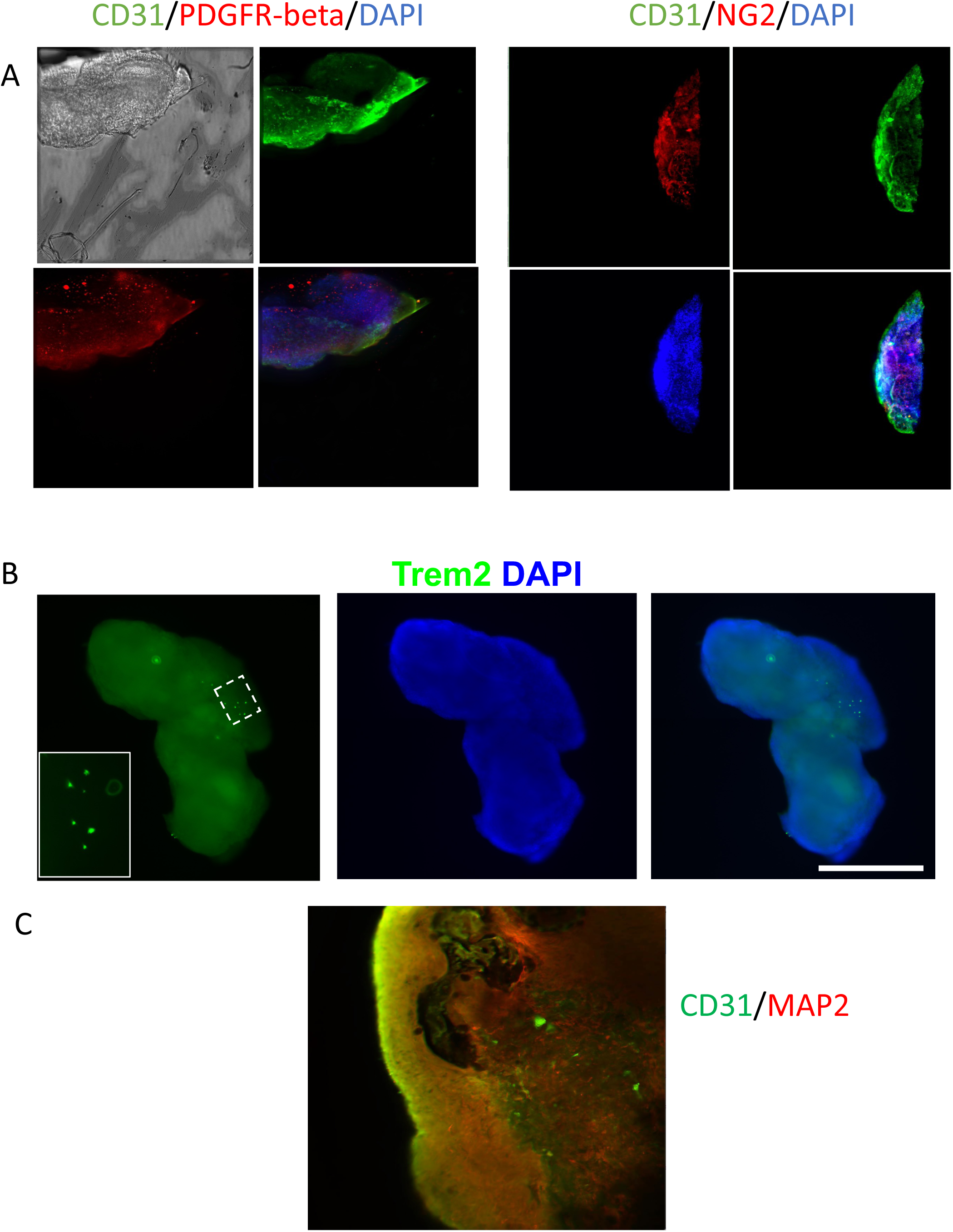
Expression of cell type specific markers in the organoids. (A), after 10 days of neuronal differentiation, pericyte markers PDGFR-beta and NG2 were expressed along with endothelial marker CD31. (B), Microglial marker Trem2 was expressed in the organoid. (C), Vascular-like structures developed in later mature stage of organoids that immunostained with endothelial marker CD31 and neuronal marker MAP2.

### ScRNA-Seq analysis showed that the organoids contain neurons and glia like in human brains

Four organoids were dissociated separately for scRNA-Seq analysis. The results showed that as expected, they were built with neurons, oligodendrocytes, microglia, astroglia and decent amount of neural stem cells and pluripotent stem cells (Fig. 3A). A pseudo development trajectory (Fig 3B) showed how these cells were differentiated from pluripotent stem cells over the time. When compared the organoids with the data from human brain samples obtained from Allen Brain Atlas, the organoids contained varieties of neurons and glia comparable to the types found in human brains (Fig 4).

**Figure 3.**
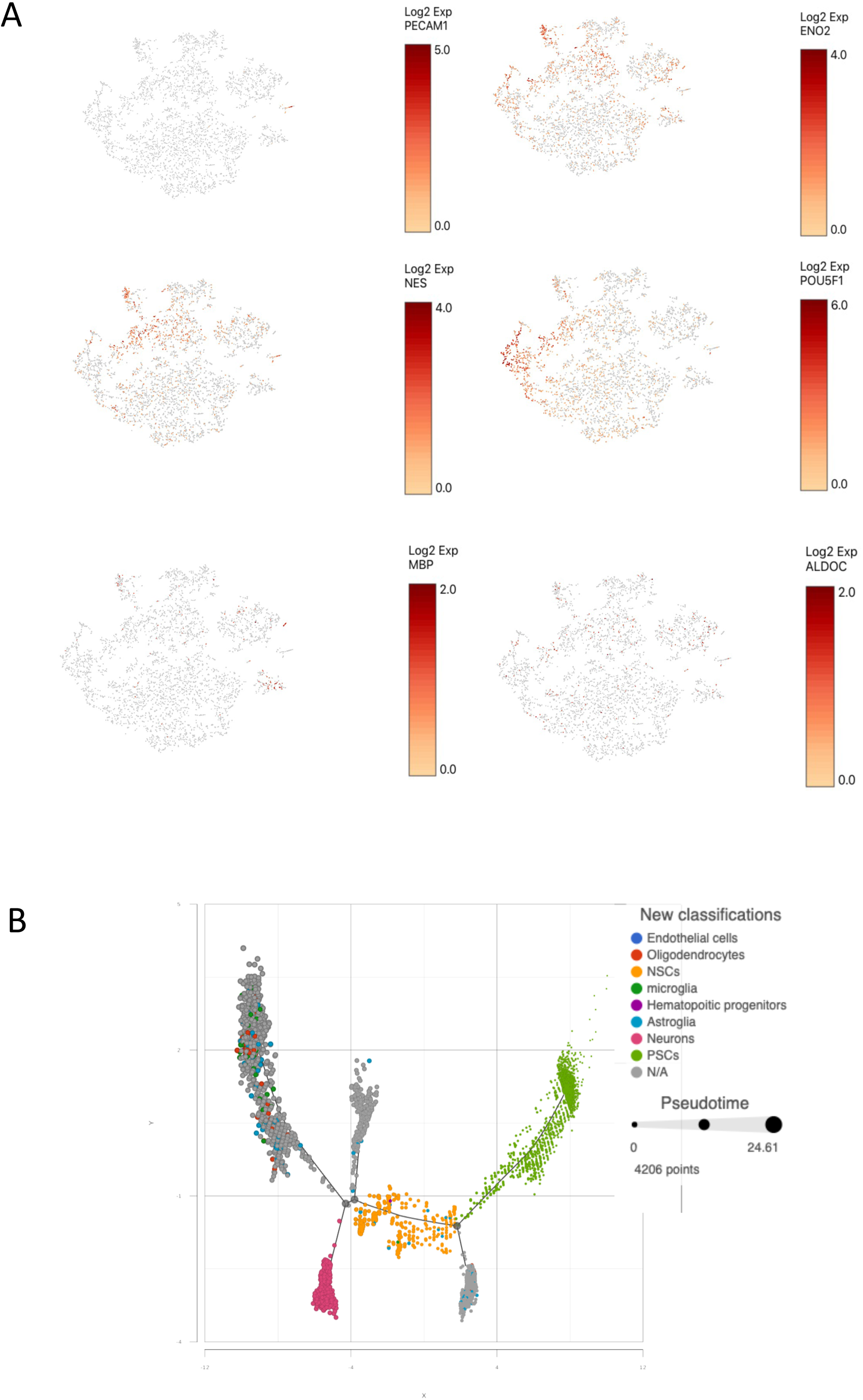
ScRNA-seq analysis of the cellular development of organoids. (A), the transcripts of cell type specific markers in the brain organoids were studied using scRNA-Seq analysis and t-SNE plots showing the organoids were consisted of cells expressing cell type specific markers such as for endothelial cells (PECAM1), neurons (ENO2), neural stem cells (NES), pluripotent stem cells (POU5F1), oligodendrocytes (MBP) and glial (ALDOC) cells. (B), a pseud time graph showing the trajectory development of the cell types within the organoids.

**Figure 4.**
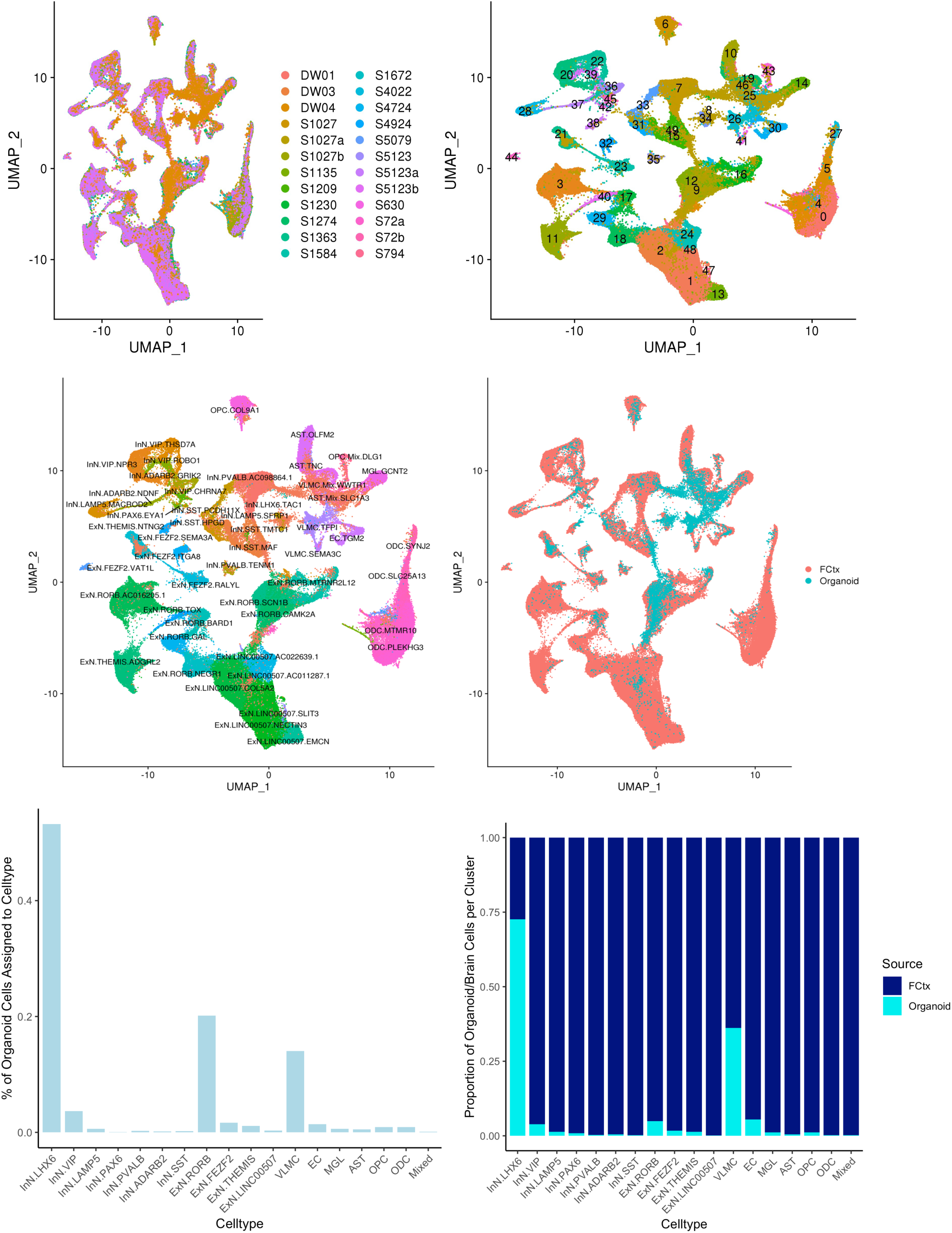
Comparison cell types within the organoids and human brains. ScRNA-Seq data from the organoids and human cerebral brain tissues were compared and presented using UMAP and assigned to specific cell types.

### Gene profiling in EC-neural organoids emphasized functions of angiogenesis and vascularization

We thus named the organoids containing both endothelial cells and other neural cells as EC-neural organoids. To compare the difference between EC-neural organoids and the classical brain organoids which lack endothelial cells, we derived both EC-neural organoids and traditional cerebral organoids from six iPSC lines side by side. At four weeks old, the organoids were collected for gene expression profiling using RNA-Seq analysis. To test for and select differential expressed genes, the raw expression per gene were first pedestalled in scaled TPM units by adding a value of 2 before undergoing the Log2 transform. Then the cyclic lowess normalization was applied to correct for differences in expression distribution spread and location. Where after, the mean expression was modeled by the coefficient of variation to identify at what expression value the coefficient of variation exceeded 20% = value = 5. The genes expression less than this value were then filter removed and any values less than this value was floored to this value. When selection of genes to be differential between N vs E if they had an absolute linear fold change > 2X and an uncorrected p-value < 0.01, this provided for a set of 62 genes. As summarized in the volcano plot (Figure 5A). The covariance-based PCA is regenerated using only the differential genes selected and also the companion clustered heatmap (SFig 2A and B). In addition, applied WGCNA (=Whole Genome Correlation Network Analysis) was also performed in attempt to find modules of genes that have expression that both covaries with one another that also has correlation with the class of interest = N vs E. The result is shown in the summary heat map that describes the modules identified (y-axis) and their correlation (p-value) to N vs E (SFig 2C). While there are no modules that have a (p-value) < 0.05, the closest two modules that come close are the turquoise module and the grey module. While genes in the turquoise module have lower expression in N than E, the genes in the grey module have higher expression in N than E. The top expressed genes in turquoise modules was IGFBP6 and in grey module was KRBA2, as shown in SFig 2D. The significant differential gene expressions were also confirmed by RT-PCR analysis (Fig 5E). The top 20 enriched pathways and functions were illustrated as Figure 5C and 5D.

**Figure 5.**
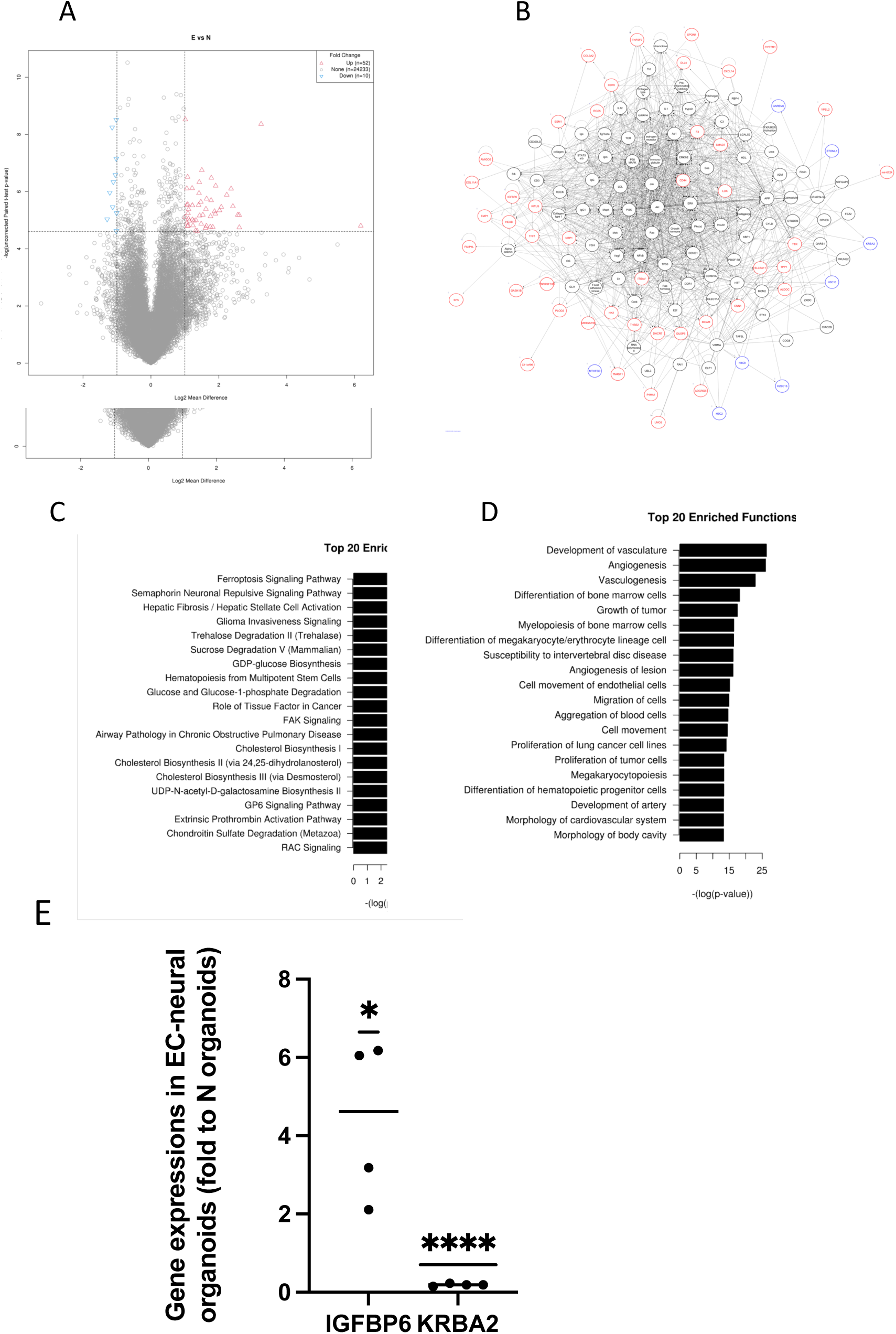
RNA-Seq analysis revealed the difference between EC-neural organoids and cerebral organoids. At 4 weeks old, EC-neural organoids (E) and traditional cerebral organoids (N) generated from 6 iPSC lines were collected for RNA-Seq analysis. Differences of gene expression were observed (A) and their relations were plotted as in (B). The top 20 enriched pathways (C) and functions (D) were presented. (E), The difference of gene expressions of IGFBP6 and KRBA2 between E and N were also confirmed using RT-PCR N=4, * P<0.05. **** P<0.0001.

Those differential genes between E vs N condition that have connectivity with the top enriched pathway (Ferroptosis Signaling Pathway) and the top enriched functions (Development of vasculature, Angiogenesis, Vasculogenesis) as well as the other genesis-related functions were plotted in networks as in Figure 5B. The nodes are colored by linear fold change (blue = down-regulated, red = up-regulated) Given all but one are down-regulated, it would seem the E condition would in turn be more “pro” outcome than N condition per Development of vasculature, Angiogenesis, and Vasculogenesis.

### KRBA2 is a critical factor regulating neuronal differentiation

As expressions of IGFBP6 and KRBA2 showed significant difference between EC-neural organoids and neuronal only organoids, we used RNAi treatment to knock down the genes in neural stem cells and determined their effects on a variety of genes involving neuronal differentiation. Although both RNAi knocked down the corresponding genes significantly, the IGFBP6 inhibition showed no significant effect on the genes we checked in the neural stem cells. On the contrary, the inhibition of KRBA2 resulted in significantly lower expression of nestin and MAP2, markers for neuronal differentiation but no significant effect on OCT4 and HERV-K Env, genes associated with pluripotent stemness (Fig 6). These results indicating the EC-neural organoids could be used as a tool to study pathways for neuronal specific developments.

**Figure 6.**
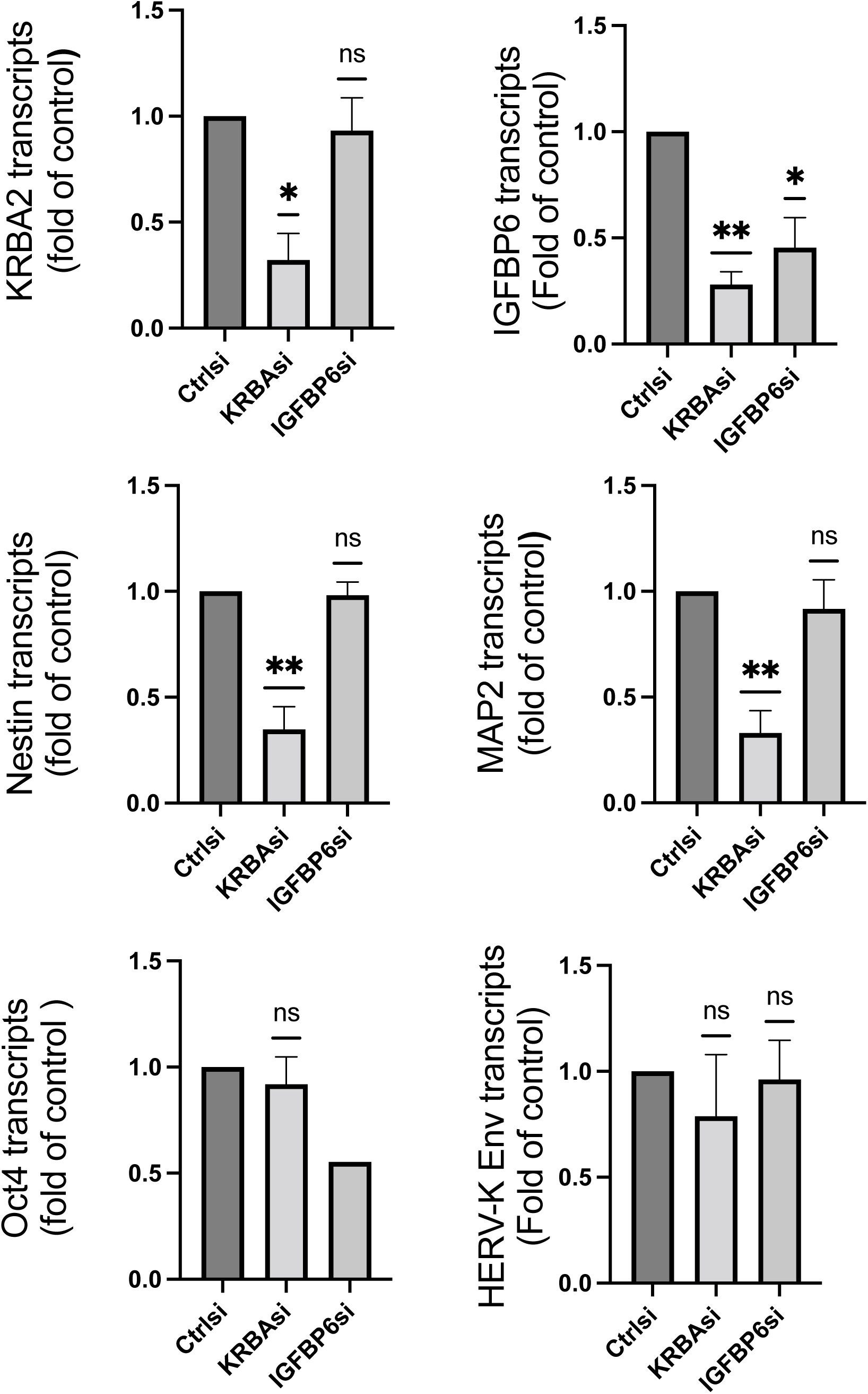
KRBA2 knockout decreased the expression of markers for neuronal differentiation. Neural stem cells were transfected with siRNAs targeting KRBA2 or IGFBP6. The total RNAs were extracted after 48 hours for RT-PCR study. The knockout of KRBA2 significantly inhibited gene expressions of neuronal markers Nestin, MAP2 and IGFBP6 but not markers for stemness OCT4 and HERV-K Env. N=4. * P<0.05, **P<0.01, compared with negative control siRNA.

### EC-neural organoids showed low level of ACE2 and TMPRSS2 expressions comparable to published databases for SARS-Cov-2 receptors

We took advantage of published human brain databases to determine the expression of certain factors which play important roles in mediating SARS-CoV-2 infectivity. Using Allen’s Atlas microarray database(15) (http://human.brain-map.org), it was found that expression of ACE2 and TMPRSS2 were low across the different brain regions, while FURIN and BSG expressions were relatively higher (SFig 5). To determine the possible cell types that may be infected by SARS-CoV-2, we used the cell transcriptomic RNA-Seq data tool in the Allen brain database(15), which confirmed that there was no significant expression of ACE2 or TMPRSS2 in neural cells (SFig 5) (http://celltypes.brain-map.org/rnaseq/human_m1_10x). The similar results were also found in another adult human brain RNA-Seq dataset(18) (http://www.gbmseq.org) (SFig 6).

We checked the gene expressions of SARS-Cov-2 receptors in the EC-neural organoids. Results showed that ACE2 or TMPRSS2 were expressed in a few numbers of cells in the organoids. However, there were more cells with alternative SARS-CoV-2 binding proteins Furin, NRP1 and BSG. (SFig 1). Both glia and neurons express ACE2 and TMPRSS2, though the percentage of positive cells was relatively higher in glial cells compared to neurons (SFig. 4). These results agree with the results we observed using on-line scRNA-Seq databases on human brain. These data suggest an alternative mechanism of virus infection in the brain, if any, compared to other organs.

## DISCUSSION

The generation of endothelial cells and blood vessels has been challenging in iPSC - differentiated neural organoids as endothelial cells and neural cells are developed from different germ layers. While most neural cells are originated from ectoderm, endothelial cells are given rise from mesoderm. The traditional protocols for neural organoids involve neural induction as soon as embryoid bodies are formed, which determines the ectoderm fate, making differentiating cell types to other germ layers almost impossible. Thus, we decided to approach in a way similar to in vivo fetal brain development by serially induce endothelial and neural differentiation. This approach turned out to be successful as not only neurons and endothelial cells were presented in the organoid, other cell types such as microglia could also thrive.

The blood vascular development consists of the initial vasculogenesis and then angiogenesis. The vasculogenesis is when the angioblasts differentiate into endothelial cells and give rise to the de novo vascular networks(19). The necessity of human brain vascular system development begins when the neural tube is isolated from the surrounding amniotic fluid by dense connective tissue and the meninx primitive. From the later, the primitive vascular loops develop and form the “meningeal meshwork”, within which preferential streams appear and raise to the subarachnoidal brain arteries (20). The typical brain angiogenesis thus is characterized by proliferation of endothelial cells which penetrate the neural tissue from the brain surface to form interconnected cannels. Based on this knowledge, we mimicked our protocol by starting endothelial cell induction first. By starting endothelial cells induction using BMP-4, bFGF plus Matrigel treatment, we provided a relatively closed environment for the organoid development, where the outlayer of cells are induced to form PECAM-1 positive endothelial cells, which then proliferate preferably on the Matrigel coating layer to make a cyst that prevent the inner cells for fully interact with the outside factors, allowing them to differentiate to other cell types such as neural cells. It is reported that endothelial cells with stem cell properties exist as CD157 (also known as Bst1) positive cells. The CD157+ endothelial cells contribute to endothelial cell turnover and capable of reconstitute hierarchical blood vessel networks in vivo(21). We observed abundant CD157+ cells in the organoids when using scRNA-Seq analysis, indicating there are endothelial stem cells and the potential of the organoids to maintain its vascular system.

We observed in the EC-neural organoids that the endothelial cells are first developed lining on the surface, covered and isolate the inner mass of the neural tissue. The endothelial cells would only penetrate the neural tissue at later time points when the organoids reached ages beyond 8-12 weeks. We also observed that connective tissue markers within substantial number of cells in the organoids. These results indicate that in the organoids, the endothelial cells and vascular like structures develop in a similar way as observed in human brain, by isolating the neural tissue with connective tissue and a meningeal surface first then penetrating from the surface into the neural tissue mass. As a result, the vascular cannel structures in the brain only happen in a later time as well. It is reasonable to believe that the angiogenesis within the organoids is also a dynamic process which would continue to develop along with the organoid age.

However, we should point out that the chances of tissues from other germ layers /organs may also rise within the organoids, likely due to the early enclosure of the cell mass from the immediate environment, which prevent direct perfusions and subdue the effect of directional differentiation. In fact, we have observed AFP (alpha fetoprotein) in the early organoids which then decrease along the time, likely because of neuronal selective media. We also observed other no-specific markers in the organoids using the RNA-Seq analysis. It is possible that additional organoid types or even with a structure consisting with multiple organoid identities can be developed by manipulating the directional signals after the initial endothelial cells’ formation and Matrigel enclosure.

To explore the potentials of using the organoids to study neuronal development, we compared the gene expression profiles between 3D organoids comprising endothelial cells and without endothelial cells. The top differential pathways are on endothelial cells and blood vessel differentiation and development. IGFBP6 expression is significantly higher in the ones with endothelial cells, while KRBA2 is higher in organoids without endothelial cells. IGFBP6 has been reported being expressed by endothelial cells(22) and playing important roles in blood vascular development such as maintaining endothelium integrity(23) and mediating vascular smooth muscle cell proliferation and morphologies (24), It has been reported that serum IGFBP6 level is higher in young mice compared to aged ones(25). It is secreted by various stem cells and blasts together with TIMP2(26–28), a factor playing key role in revitalizing aging hippocampal function(29). Published reports suggested that IGFBP6 may also play some roles in the process of preserving cells against aging effects, such as resisting the cell senescence(30) and alleviating neurodegenerative disorders including PD(31). Thus, endothelial cells by secreting IGFBPs, could help reconstitute brain hemopoiesis. We found the upregulated IGFBP6 expression in EC-organoids compared to neural-only organoids, also proved the important roles of IGFBP6 could play in the brain development and function, which could not be studied in a traditional generated brain organoid.

We found there was higher level of KRBA2 expression in the neural only organoids. When the KRBA2 expression was knocked down by siRNA in neural stem cells, we observe a general depression of gene expression, except on OCT3/4, indicating KRBA2 may play an important role in regulating neural differentiation. KRBA2 was thought a result of horizontal transfer of GINGER2 transposon from insect to human genome, The function of KRBA2 is unclear, although the similar KRAB domain has been reported play an repressive role in gene regulation by recruiting transcriptive repressor complexes(32). It has been suggested that KRBA2 may be adopted by hosts playing a defensive role against other mobile elements. Given that retroelement activities are usually higher in stem cells but decrease during differentiation, KRBA2 may indeed function as a transcriptional factor facilitating neural differentiation given its broad band enhancing effect on neuronal specific gene expressions.

We also examined the SARS-CoV-2 receptors on the EC-neural organoids to see if it be suitable for study of the virus infection. The COVID-19 pandemic has had an unprecedented impact on human life. Besides the direct effect of SARS-CoV-2 on organs such as lung, kidney and cardiovascular system, there is increasing concern about the involvement of central nervous system. While neurological manifestations are commonly seen in patients with the infection, they can also be the presenting manifestation in some(33). Yet, it is unclear if the virus can directly invade the brain. In rare cases, SARS-CoV-2 has been demonstrated in the cerebrospinal fluid in patients with meningo-encephalitis (34, 35) and in the brain at autopsy in endothelial cells by electron microscopy(36, 37). In another patient, the virus was demonstrated budding from myelin sheaths in the medulla and olfactory pathways(38). However, others have been unable to detect the virus in the CSF or brain at autopsy(39).

We took advantage of published human brain databases to determine the level of expressions of genes that may mediate SARS-CoV-2 infection. Using Allen Atlas’s microarray database, we found both ACE2 and TMPRSS2 expression were low across the different brain regions, while the expression of FURIN and BSG were relatively higher. To determine the possible cell types that may be infected by SARS-CoV-2, we used the scRNA-Seq data tool in Allen Brain database, which confirmed there was no significant expression of ACE2 or TMPRSS2 in neural cells. The lack of TMPRSS2 expression and low level of ACE2 expression in neural cells was confirmed in another brain scRNA-Seq dataset from adult brain samples from UCSF. In looking at developing human brain sc-RNA-Seq data, we found low levels of expression of both ACE2 and TMPRSS2. To further explore these findings, we investigated the expression of these genes using the newly developed 3D brain organoid model with incorporation of endothelial cells.

A recent study used 3D brain organoids to study the infectivity of SARS-CoV-2 in the brain and found that the virus was capable of infecting the organoids (40). However, this model of brain organoids lacked endothelial cells or a blood-brain-barrier (BBB). In the absence of the blood brain barrier, the neuronal cells would be exposed to very high concentrations of the virus which would not mimic the in vivo system. Besides, the vascular endothelium has been reported to be the primary target of SARS-CoV-2 (36) and high level of ACE2 is expressed in brain vascular pericytes in mouse brain(41), indicating an important role in neuropathogenesis of the infection. The brain organoids with differentiated endothelial cells, which initially covers the organoids on the surface but later penetrates into the mass to make vessel like structures. It is highly consistent with the process during embryonic development, when the perineural vascular plexus covers the developing neural tube into which vessels subsequently penetrate (42). These unique properties make a BBB-like membrane that separates the neural cells from the external environment and thus more closely resembles in vivo conditions, although the vessels lack blood circulation. We found that in the brain organoids, there are only a few cells that showed low levels of ACE2 or TMPRSS2 expression. There is no ACE2 or TMPRSS2 expression in CD31+ (PECAM) endothelial cells or PDGFRβ+ CD146+ NG2+ pericytes. This lack of ACE2 expression in pericytes may be due to the different species as the previous published results were mainly done in mouse models. ACE2 was higher in glial cells including MBP expressing oligodendrocytes and was present in very few neurons. However, ACE2 and TMPRSS2 were not co-expressed in these cells, indicating that the brain cells may not be initial targets of the virus. This still raises the possibility that the brain could be infected via the ACE2 receptor if the cells were exposed to high levels of the virus upon disruption of the blood brain barrier. Viral transmission could possibly also occur via cell-to-cell contact and could spread along neuronal pathways. However, these possibilities need to be further explored.

It is possible that co-expression of ACE2 and TMPRSS2 may not be necessary for SARS-CoV-2 infection and that Furin and other such enzymes may facilitate viral entry. It has been reported that Furin, a proprotein convertase, could pre-activate SARS-CoV-2 spike protein, and thus reduce its dependence on the target cell proteases for cell entry (43). As there are high levels of furin expression in brain cells, TMPRSS2 expression may not be necessary for SARS-CoV-2 infection in the brain. However, the low levels of ACE2 are still the major limiting factor for the SARS-CoV-2 infectivity in neural cells.

As a summary, by serial induction of differentiation, we make a 3D organoid containing endothelial and neuronal cells, mimicking a vascularized developing brain. This 3D human organoids could be a useful tool to study human development involving endothelial and neuronal cell interactions and for neural infectious disorders when the blood-brain-barrier and endothelial cells are pathologically targeted.

## Supporting information

SFig

## Acknowledgments

This work is funded by NINDS intramural research funds. The authors declare no competing financial interests.

## Author Contributions

Conceptualization, T.W. and A.N.; Methodology, T.W., A.G.E., M.R.C. and A.N.; Investigation, T.W., A.B., V.M., B.D.G.,J.S., A.G.E., K.J. and R.L.; Formal Analysis: T.W., A.B, A.G.E., K.J. and R.L.; Writing – Original Draft, T.W., V.M., A.G.E. and A.N.; Writing – Review & Editing, T.W., A.B., B.D.G and A.N.; Supervision, T.W., and A.N.

## Figure legends

SFig1. <u>Expressions of SARS-CoV-2 binding associated proteins in 3D neuro-EC organoids.</u> The cell distributions of ACE2, TMPRSS2, Furin and NRP1 in four organoids were detected by scRNA-Seq analysis. The data were presented as % of SARS-CoV-2 binding protein positive cells.

SFig2. RNA-Seq analysis showed the IGFBP6 gene expression increased in EC-neural organoids (E) compared to cerebral organoids (N). While KRBA2 expression increased in cerebral organoids compared to EC-neural organoids.

SFig3. Immunostaining of the organoids for endothelial cell marker CD31 and neuronal marker beat-III-tubulin.

SFig4. SARS-CoV2 infection associated gene expression in the EC-neural organoids per cell types.

SFig5. scRNA-Seq data from Allen Brain Atlas showed low levels of ACE2 and TMPRSS2 expression across brain cell types.

SFig6. scRNA-Seq data from http://www.gbmseq.org/ showed low levels of ACE2 and TMPRSS2 expression in brain cells.

## Notes

### Competing Interest Statement

The authors have declared no competing interest.

