## Supplementary figures and images for "3D organoids containing endothelial and neural cells generation by serial inductions of differentiation on human iPSC-derived embryoid bodies"

### SFig

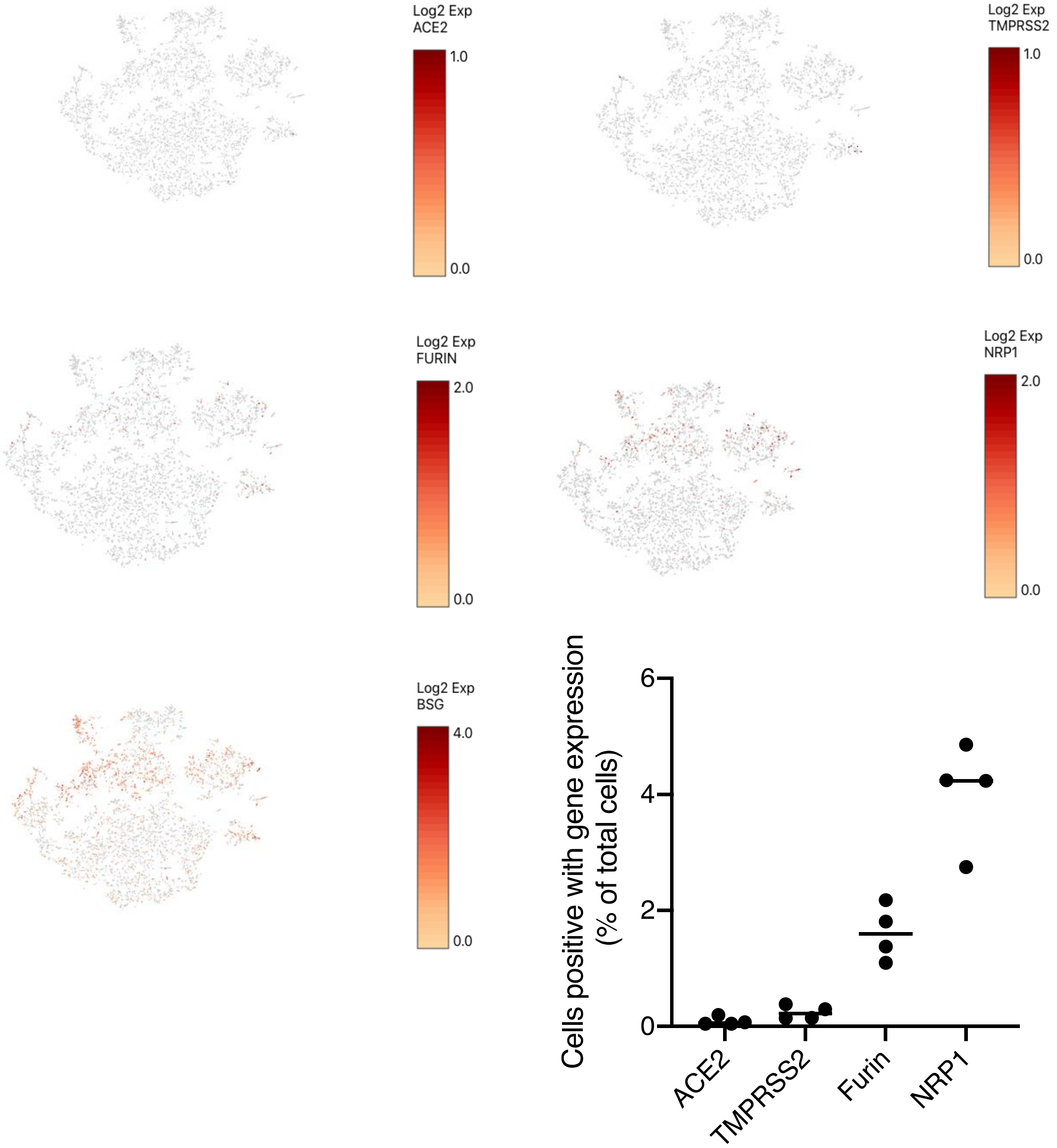

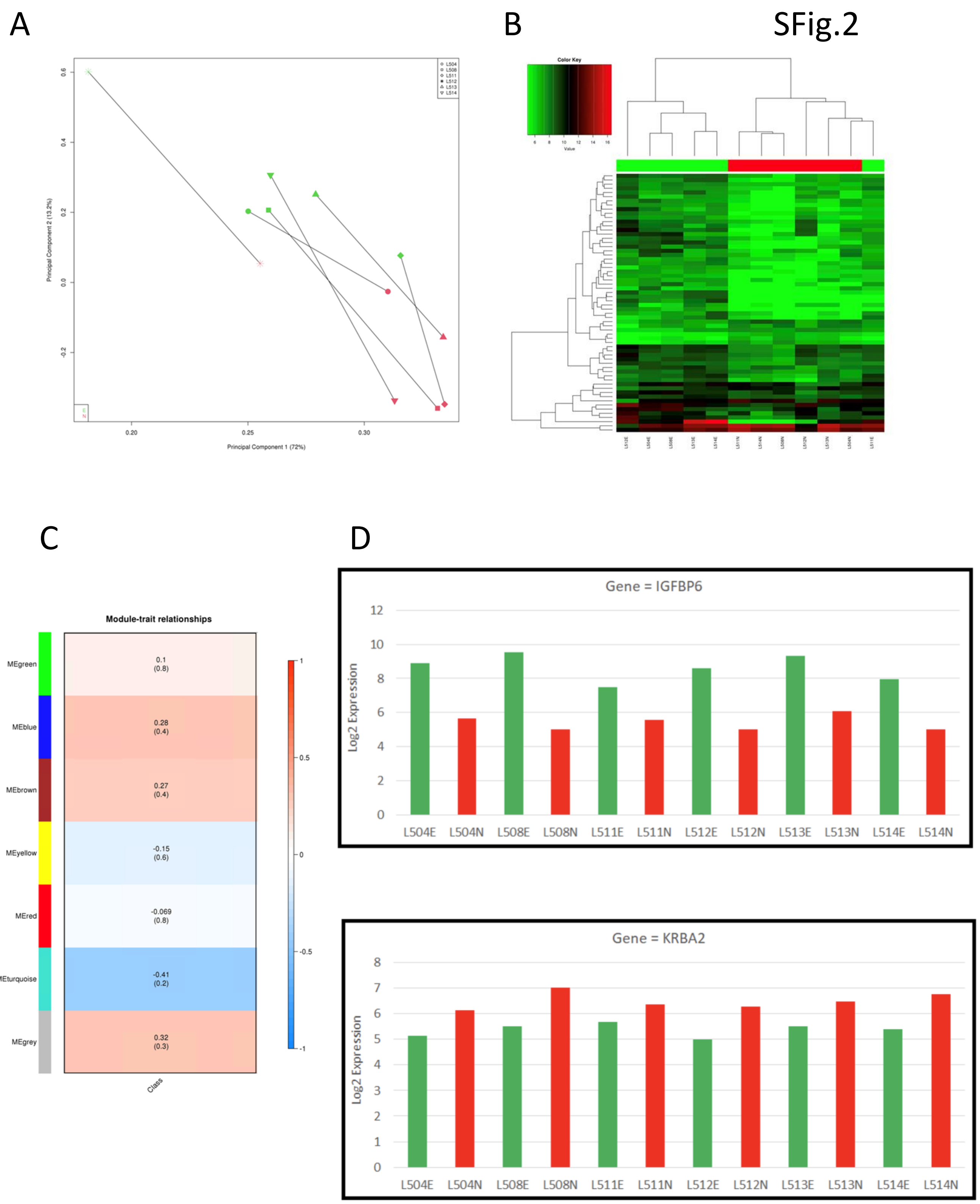

SFig.3

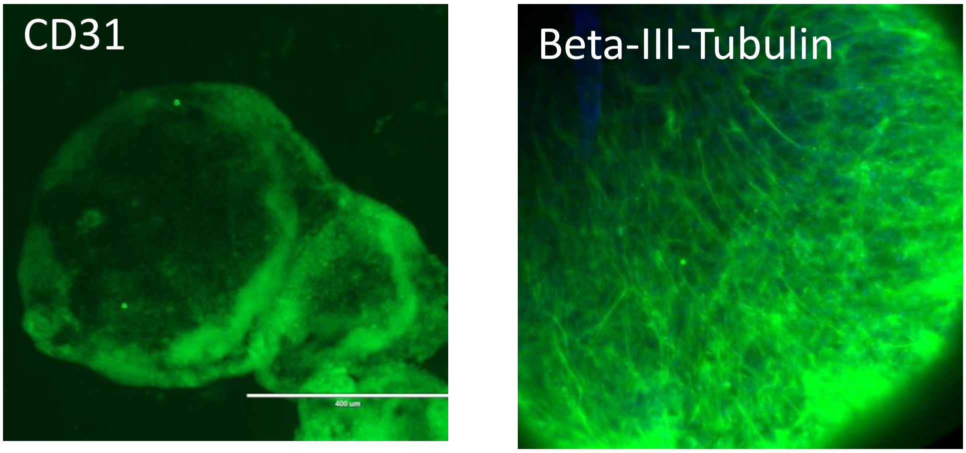

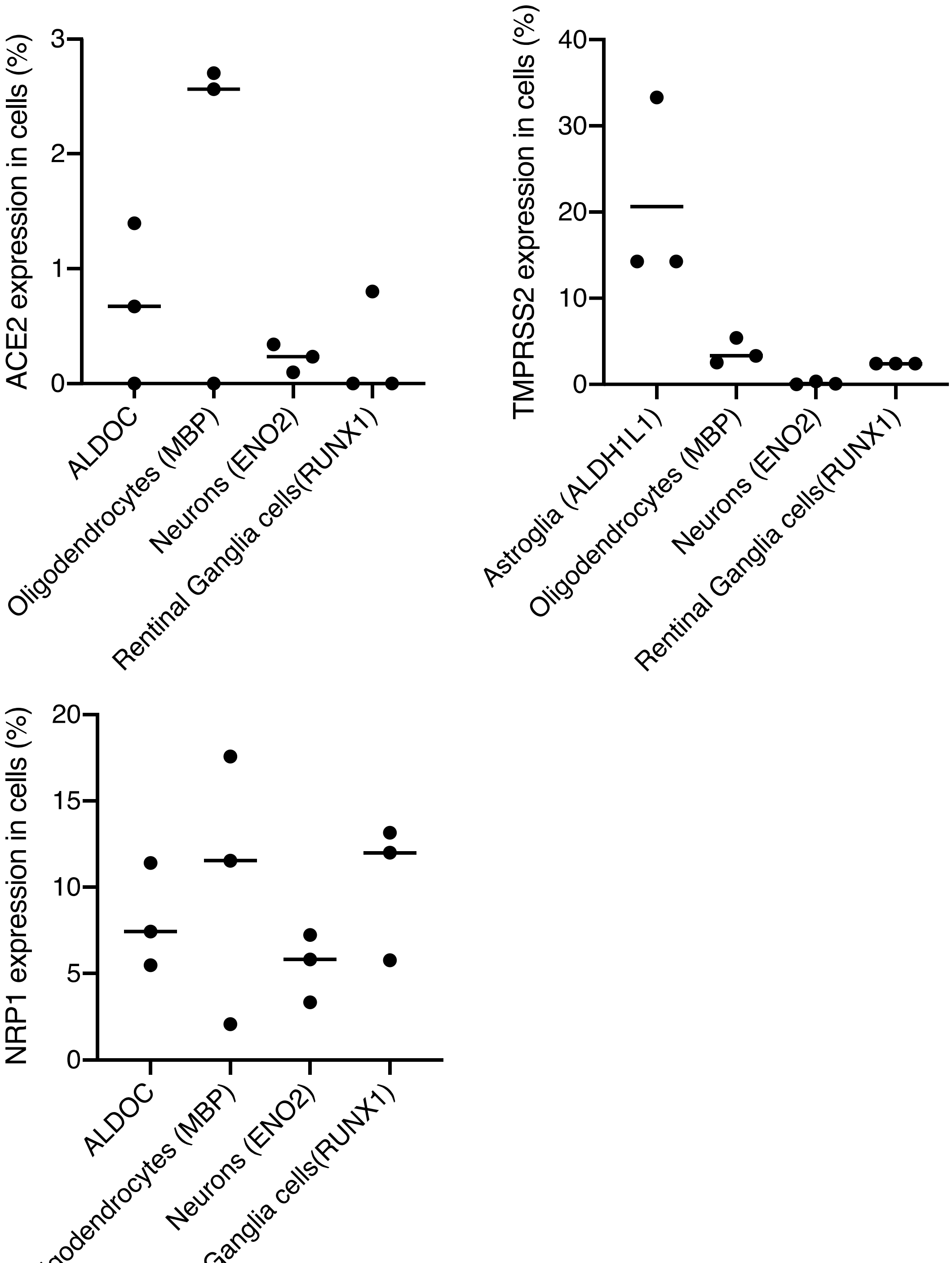

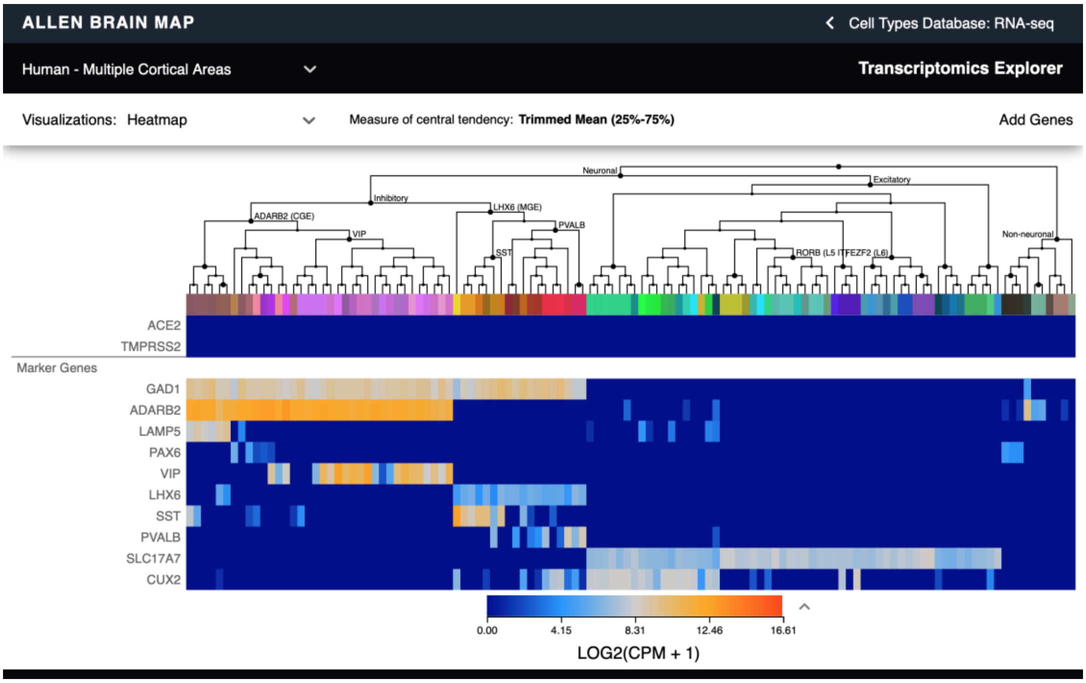

SFig.6

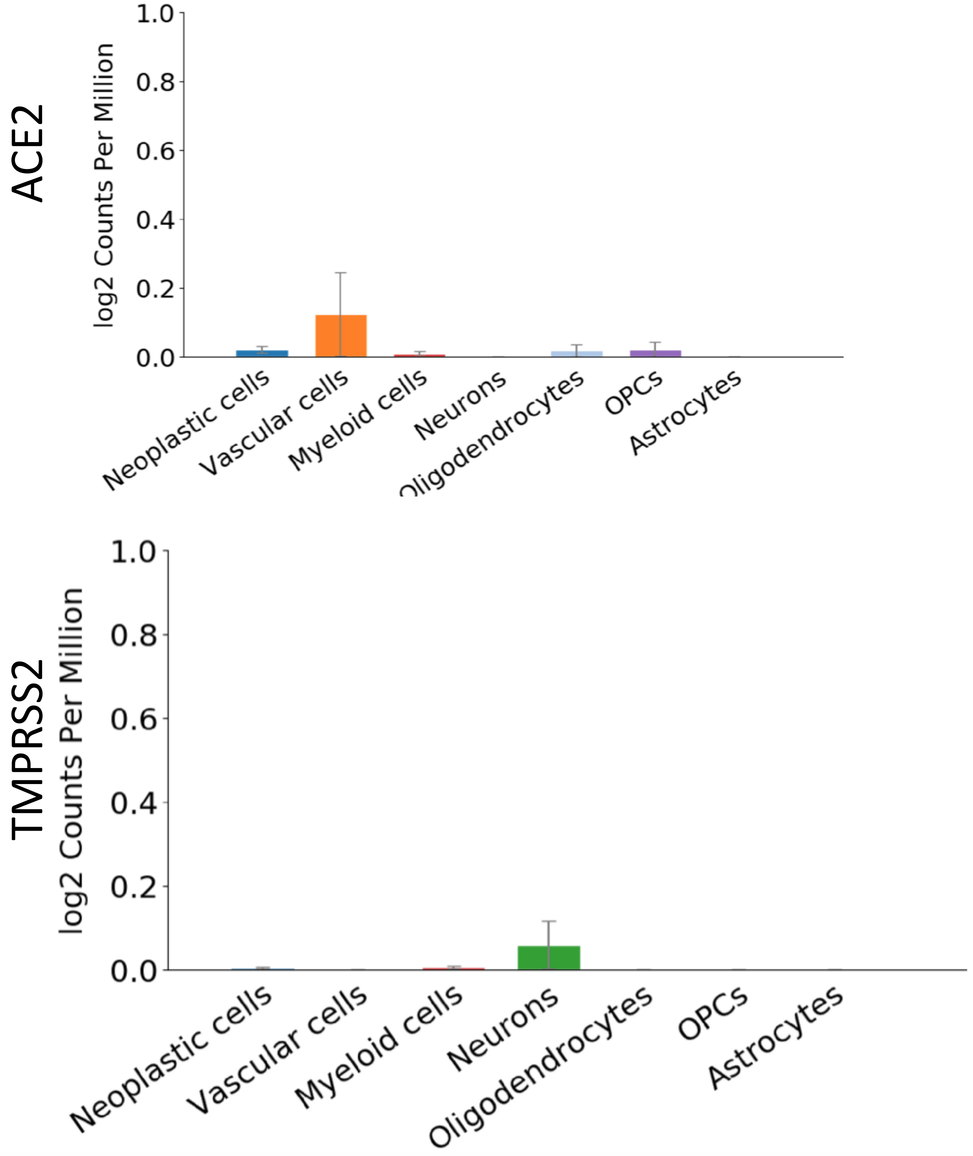


http://www.gbmseq.org/search
